# Structural basis for transfer RNA mimicry by a bacterial Y RNA

**DOI:** 10.1101/312249

**Authors:** Wei Wang, Xinguo Chen, Sandra L. Wolin, Yong Xiong

## Abstract

Noncoding Y RNAs are present in both animal cells and many bacteria. In all species examined, Y RNAs tether the Ro60 protein to an effector protein to perform various cellular functions. For example, in the bacterium *Deinococcus radiodurans*, Y RNA tethers Ro60 to the exoribonuclease polynucleotide phosphorylase, specializing this nuclease for structured RNA degradation. Recently, a new Y RNA subfamily was identified in bacteria. Bioinformatic analyses of these YrlA (Y RNA-like A) RNAs predict that the effector-binding domain resembles tRNA. We present the structure of this domain, the overall folding of which is strikingly similar to canonical tRNAs. The tertiary interactions that are responsible for stabilizing tRNA are present in YrlA, making it a close tRNA mimic. However, YrlA lacks a free CCA end and contains a kink in the stem corresponding to the anticodon stem. Since nucleotides in the D and T stems are conserved among YrlAs, they may be an interaction site for an unknown factor. Our experiments identify YrlA RNAs as a new class of tRNA mimics.

## 1 INTRODUCTION

In addition to the canonical tRNAs that function in protein synthesis, several RNAs depend on structural similarity to tRNA in order to function. These tRNA mimics include the bacterial transfer-messenger RNA (tmRNA) that rescues stalled ribosomes from mRNAs lacking stop codons and the tRNA-like structures that contribute to translation and replication of positive-strand RNA viruses. For tmRNA, the tRNA-like portion undergoes aminoacylation and binds the elongation factor EF-Tu, allowing it to enter the A-site of arrested ribosomes and function as an acceptor for the stalled polypeptide (Keiler, 2015). Although the viral tRNA-like sequences have diverse functions in translation and replication, several are substrates for the CCA-adding enzyme and a tRNA synthetase and interact with the elongation factor EF-1A (Dreher, 2009). Additionally, tRNA-like structures at the 3′ end of several mammalian long noncoding RNAs (lncRNAs) are cleaved by the tRNA 5′ maturation enzyme RNase P, resulting in 3′ end formation of the upstream lncRNA and release of a tRNA-like noncoding RNA (ncRNA) of unknown function (Sunwoo et al., 2009; Wilusz et al., 2008).

Another class of RNAs that are proposed to mimic tRNA consists of bacterial ncRNAs known as YrlA (Y RNA-like A) RNAs (Chen et al., 2014). These ncRNAs are members of the Y RNA family, 80 to ~220 nucleotides (nt) ncRNAs that were initially identified in human cells because they are bound by the Ro60 autoantigen, a major target of autoantibodies in patients with systemic lupus erythematosus (Wolin et al., 2013). Studies in vertebrate cells revealed that Y RNAs regulate the subcellular location of Ro60 and its association with other proteins and RNAs. For example, Y RNA binding masks a nuclear accumulation signal on Ro60, retaining it in the cytoplasm (Sim et al., 2009). Y RNAs also scaffold the association of Ro60 with other proteins, as binding of the zipcode-binding protein ZBP1 to a mouse Y RNA adapts the Ro60/Y RNA complex for nuclear export (Sim et al., 2012). Moreover, the ring-shaped Ro60 binds the 3′ ends of some misfolded ncRNAs in its central cavity and adjacent structured RNA regions on its outer surface (Fuchs et al., 2006; Stein et al., 2005). Because Y RNAs bind overlapping sites on the Ro60 outer surface, Y RNAs may regulate the access of misfolded ncRNAs to the Ro60 cavity (Stein et al., 2005).

Studies of bacterial Y RNAs have revealed that, as in animal cells, their functions are intertwined with that of the Ro60 protein. In *Deinococcus radiodurans*, the bacterium where Ro60 and Y RNAs have been most extensively characterized, at least two Y RNAs, called Yrn1 (Y RNA 1) and Yrn2, are bound and stabilized by the Ro60 ortholog Rsr (<u>R</u>o <u>s</u>ixty-<u>r</u>elated) (Chen et al., 2000; Chen et al., 2013; Chen et al., 2007). Consistent with co-regulation, these ncRNAs are encoded upstream of Rsr and on the same DNA strand. One role of Yrn1 is to tether Rsr to the ring-shaped 3′ to 5′ exoribonuclease polynucleotide phosphorylase (PNPase), forming a double-ringed RNA degradation machine called RYPER (Ro60/Y RNA/PNPase Endonuclease RNP) (Chen et al., 2013). In RYPER, single-stranded RNA threads from the Rsr ring into the PNPase cavity for degradation, rendering PNPase more effective in degrading structured RNA (Chen et al., 2013). In addition to its role in RYPER, Rsr assists 23S rRNA maturation by two 3′ to 5′ exoribonucleases, RNase II and RNase PH, during heat stress, where Rsr functions as a free protein, and is inactive when bound to Y RNA (Chen et al., 2007). Thus, in addition to acting as a tether, Yrn1 may function as a gate to block access of other RNAs to Rsr.

Y RNAs are modular, a feature that is critical for carrying out their functions. All characterized Y RNAs contain a long stem, formed by base-pairing the 5′ and 3′ ends of the RNA, that contains the Ro60 binding site. Although in both metazoans and *D. radiodurans*, the sequences required for Ro60 binding map to a conserved helix (Chen et al., 2013; Green et al., 1998), a structure of a *Xenopus laevis* (*X. laevis*) Ro60/Y RNA complex revealed that Ro60 primarily interacts with the 5′ strand (Stein et al., 2005). Consistent with the idea that base-specific interactions with this strand are critical for Ro60 recognition, only the 5′ strand of the helix is conserved across bacterial species (Chen et al., 2014). The other end of all Y RNAs consists of internal loops and stem-loops that interact with other proteins. For example, to form the mammalian Ro60/Y RNA/ZBP1 complex, ZBP1 interacts with the large internal loop of the Y RNA (Köhn et al., 2010; Sim et al., 2012), while in *D. radiodurans* RYPER, this portion of Yrn1 interacts with the KH and S1 single stranded RNA-binding domains of PNPase (Chen et al., 2013). Thus, one role of this second Y RNA module is to tether Ro60 to an effector protein.

Remarkably, for many bacterial Y RNAs, the effector-binding module bears a striking resemblance to tRNA. The first member of this Y RNA subfamily was identified in the enteric bacterium *Salmonella enterica* serovar Typhimurium (*S. Typhimurium*), where it and a second Y RNA were bound by Rsr and encoded 3′ to this protein (Chen et al., 2013). Because the two Y RNAs in S. Typhimurium appeared to represent a separate evolutionary lineage from the more metazoan-like Y RNAs characterized in *D. radiodurans*, these RNAs were designated YrlA and YrlB (Y RNA-like A and B)(Chen et al., 2013). Homology searches revealed that RNAs resembling YrlA were widespread, as they were detected near Rsr in >250 bacterial species representing at least 10 distinct phyla (Chen et al., 2014). Identification of conserved sequences and secondary structures within these RNAs revealed similarities to the D, T and acceptor stem-loops of tRNA (Figure 1) (Chen et al., 2014). Consistent with a tRNA-like fold, *S*. Typhimurium YrlA is a substrate for two tRNA modification enzymes, TruB and DusA, that modify the T and D loops, respectively (Chen et al., 2014).

**Figure 1.**
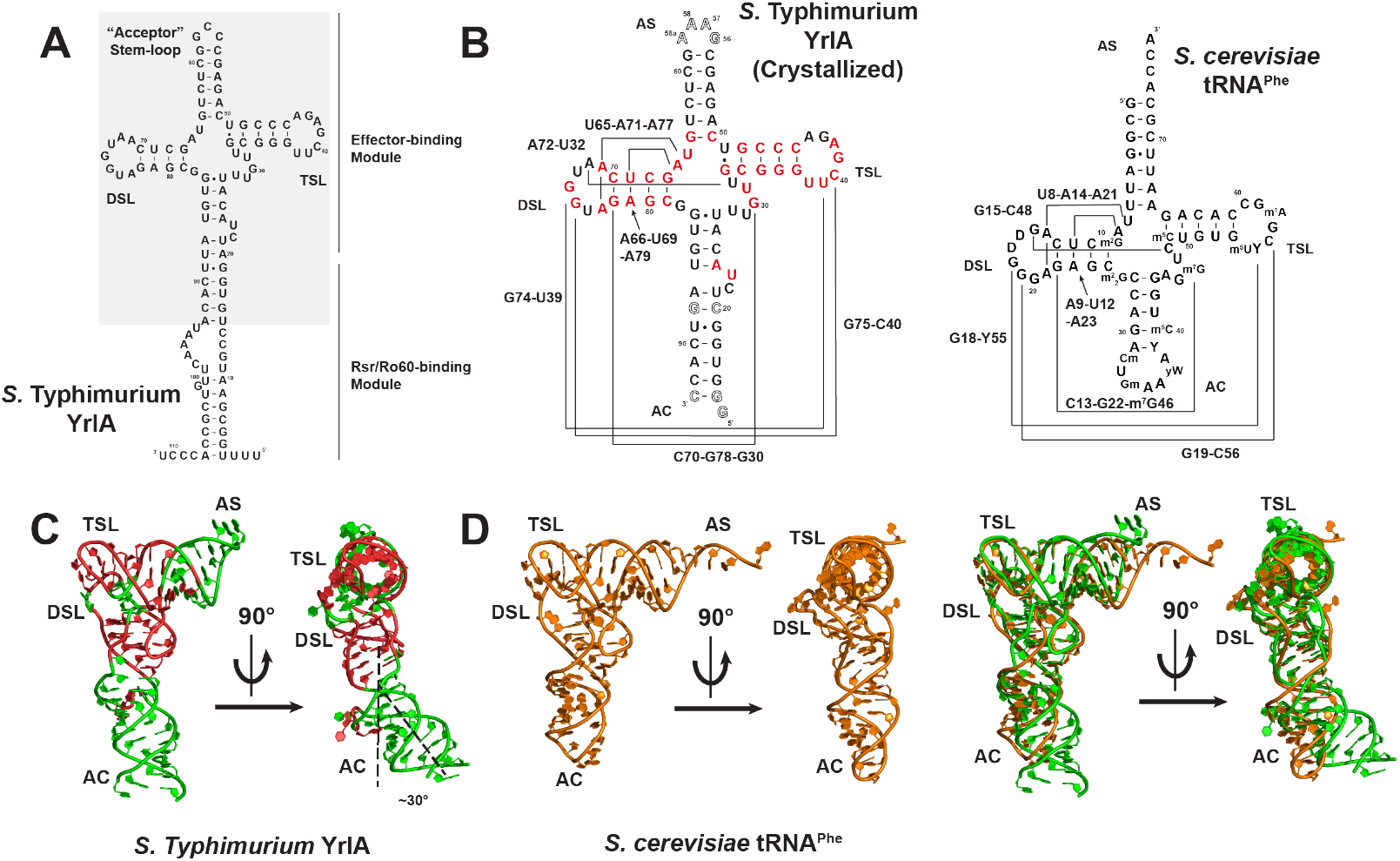
*S*. Typhimurium YrlA folds into a structure very similar to that of canonical tRNA. (A) Full-length *S*. Typhimurium YrlA consists of an Rsr/Ro60-binding module and a effector-binding module (shaded) that resembles tRNA. (B) Secondary structures of the crystallized *S*. Typhimurium YrlA effector-binding module (nt 1593) and *S. cerevisiae* tRNA^Phe^. Important tertiary interactions are connected by lines and are labeled. Conserved nucleotides in YrlA are red. Modified nucleotide sequences to facilitate crystallization are shown as hollow characters. (C) The crystal structure of *S*. Typhimurium YrlA (green, PDB: 6cu1). Conserved nucleotides in YrlA are red. (D) Left, the tertiary structure of *S. cerevisiae* tRNA^Phe^ (orange, PDB: 4tna); right, overlay of the YrlA (green) and tRNA^Phe^ (orange) structures.

To test the hypothesis that the YrlA effector-binding domain folds into a tRNA-like structure, we determined the structure of the tRNA-like domain of S. Typhimurium YrlA by X-ray crystallography. We show that the YrlA effector-binding domain indeed assumes a similar overall fold as tRNA and that the same tertiary interactions that stabilize tRNA are present in YrlA. In support of a critical role for the tRNA-like module, both the ability to fold into a tRNA-like structure and specific sequences within the structure are conserved in YrlA RNAs from a wide range of bacteria.

## RESULTS

We determined the crystal structure of the *S*. Typhimurium YrlA effector binding module (nucleotides 15-93) at 3 Å resolution (Figure1A, Figure S1). To obtain the high-resolution diffraction data, the 3-nucleotide (nt) loop ^56^CCG^58^ of YrlA was changed to a ^56^GAAA^58a^ tetraloop, where the extra nucleotide was numbered as 58a to maintain the original numbering of the subsequent nucleotides (Figure 1B). ^14^GG^15^ and ^94^C were added to the construct to increase transcription yield and stabilize the stem. The base pair A20-U88 was also mutated to C20-G88 for stem stabilization. The modified YrlA crystallized in the C2221 space group and diffracted to 3.0 Å resolution. The structure was determined by a combination of single wavelength anomalous dispersion and molecular replacement methods (see Materials and Methods section for more details). We were able to build all nucleotides into the electron density map and the refinement statistics are summarized in Table 1.

**Table 1.**
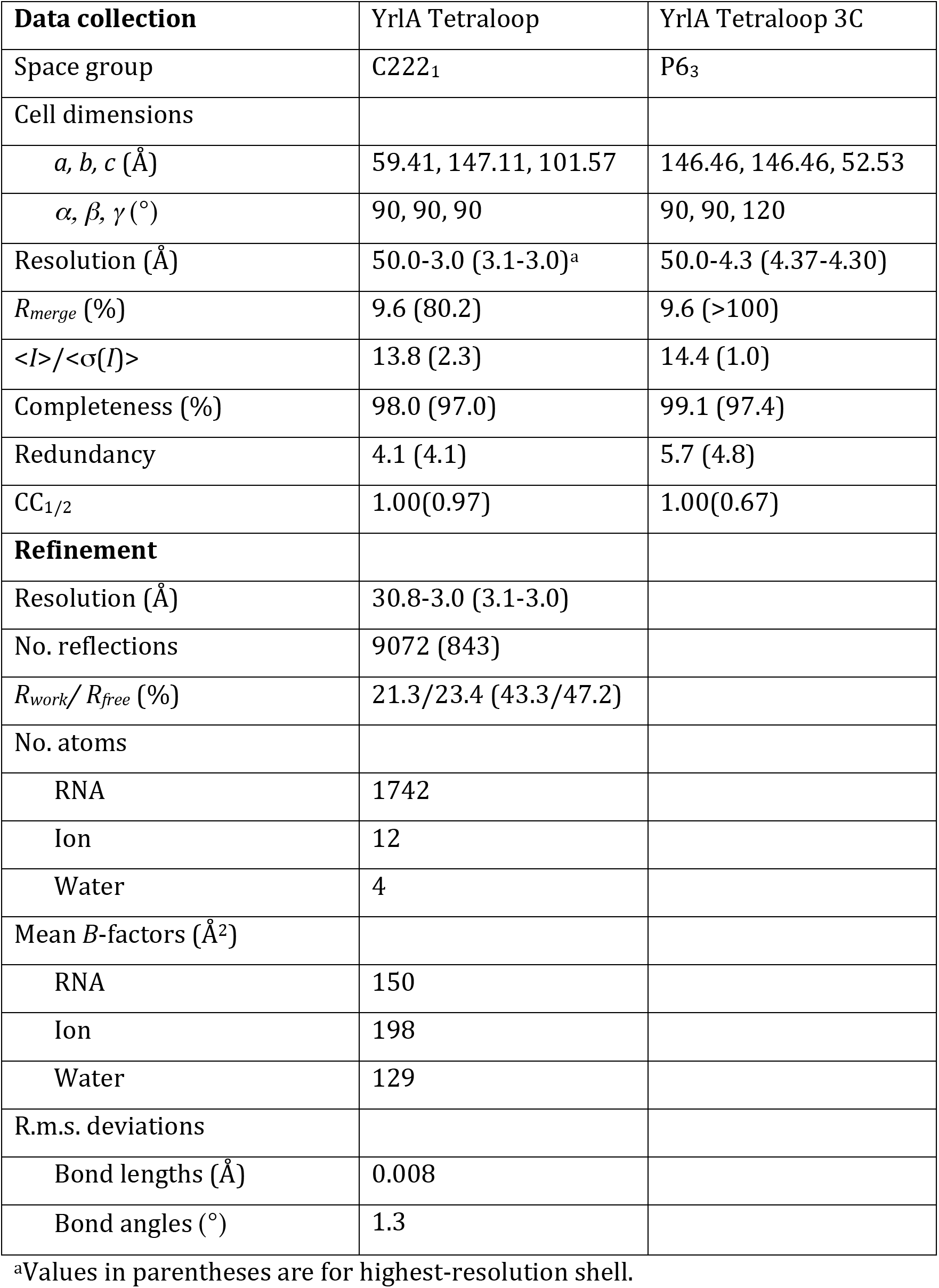
Data collection and refinement statistics (See also Figure S1)

| **Data collection** | YrlA Tetraloop | YrlA Tetraloop 3C |
| --- | --- | --- |
| Space group | C222<sub>1</sub> | P6<sub>3</sub> |
| Cell dimensions |  |  |
| *a*, *b*, *c* (Å) | 59.41, 147.11, 101.57 | 146.46, 146.46, 52.53 |
| $\alpha$, $\beta$, $\gamma$ (°) | 90, 90, 90 | 90, 90, 120 |
| Resolution (Å) | 50.0-3.0 (3.1-3.0)<sup>a</sup> | 50.0-4.3 (4.37-4.30) |
| *R*<sub>merge</sub> (%) | 9.6 (80.2) | 9.6 (>100) |
| $\langle I \rangle / \langle \sigma(I) \rangle$ | 13.8 (2.3) | 14.4 (1.0) |
| Completeness (%) | 98.0 (97.0) | 99.1 (97.4) |
| Redundancy | 4.1 (4.1) | 5.7 (4.8) |
| CC<sub>1/2</sub> | 1.00(0.97) | 1.00(0.67) |
| **Refinement** |  |  |
| Resolution (Å) | 30.8-3.0 (3.1-3.0) |  |
| No. reflections | 9072 (843) |  |
| *R*<sub>work</sub> / *R*<sub>free</sub> (%) | 21.3/23.4 (43.3/47.2) |  |
| No. atoms |  |  |
| RNA | 1742 |  |
| Ion | 12 |  |
| Water | 4 |  |
| Mean *B*-factors (Å<sup>2</sup>) |  |  |
| RNA | 150 |  |
| Ion | 198 |  |
| Water | 129 |  |
| R.m.s. deviations |  |  |
| Bond lengths (Å) | 0.008 |  |
| Bond angles (°) | 1.3 |  |

<sup>a</sup>Values in parentheses are for highest-resolution shell.

### *The overall architecture of S*. Typhimurium *YrlA resembles that of tRNA*

The YrlA RNA folds into a L-shaped structure that is characteristic of tRNAs. As predicted from the secondary structure (Chen et al., 2014), all tRNA equivalent regions are present in the YrlA structure, including a stem-loop region resembling the acceptor stem (AS, nt 50-64), the T stem-loop (TSL, nt 33-49), the D stem-loop (DSL, nt 67-81), and the anticodon stem (AC, nt 15-27 and 83-93). These structural elements fold into the L-shaped conformation with the two extended stem-loops interacting at an angle very similar to that of tRNAs (Figure 1C). The overall YrlA structure has a backbone RMSD of ~2.4 Å to that of *Saccharomyces cerevisiae* (S. *cerevisiae*) tRNA^Phe^, based on the alignment of 52 phosphorus atoms. It was shown that YrlA RNAs are substrates for tRNA modification enzymes and contain canonical tRNA modifications such as dihydrouridine in the D loop and pseudouridine in the T loop (Chen et al., 2014). The high degree of structural similarity between the two RNAs provides an explanation for the recognition of YrlAs by tRNA modification enzymes.

Deviating from the canonical tRNA structure, there is a kink in the AC region of YrlA, bending the lower portion of the stem by ~30 degrees (Figure 1C). The region extending from this stem contains the Rsr/Ro60-binding module, making YrlA much more elongated than tRNA. The two nucleotides responsible for the kink, ^23^UA^24^, are conserved among YrlA family members. However, sequence alignment indicates that these two nucleotides are predicted to be located in the variable loop (VL) of most YrlA RNAs, rather than being part of the AC stem (Chen et al., 2014). Thus, it is unclear whether the kink is a universal feature of YrlA RNAs.

A major difference between S. Typhimurium YrlA and tRNA is that the AS, which terminates in 3′-CCA in all mature tRNAs, is instead a closed loop in YrlA. (Figure 1A). In addition, YrlA lacks the anticodon loop. The length of the AS of YrlAs varies between species (Chen et al., 2014). For S. Typhimurium YrlA, the stem is only six base pairs, which is one base pair less than that of tRNAs. For other species, such as some cyanobacteria, this YrlA stem is predicted to be much longer (Chen et al., 2014). Interestingly, *Mycobacterium smegmatis* YrlA, which resembles *bona fide* tRNAs in containing a seven-base pair AS, is a substrate for RNase P. Following cleavage, the fragment corresponding to a tRNA 3′ end undergoes exonucleolytic nibbling and CCA addition (Chen et al., 2014). However, since most YrlA RNAs contain acceptor stems that are predicted to be poor RNase P substrates, the majority of YrlA RNAs likely resemble circularly permuted tRNAs with closed loop-containing acceptor stems (Chen et al., 2014).

### YrlA is stabilized by the same tertiary interactions as tRNAs

In addition to the similarities in overall folding, the tertiary interactions that stabilize the L-shaped structure of YrlA resemble those of tRNA (Figure 2). A major feature of tRNA folding is the interaction between the DSL and the TSL regions to form the tRNA elbow, which serves as a binding site for numerous enzymes that recognize tRNA (reviewed in (Zhang and Ferré-D’Amaré, 2016)). In YrlA, the interaction between nucleotides U38-A42 of the T loop and two guanines in the D loop closely mimics the elbow region of tRNA (Figure 2A, 2B). The first two nucleotides in the YrlA T loop, U38 and U39, stack with nucleotides in the T stem. The third nucleotide, C40, interacts with G75 in the DSL. There is a gap between the fourth and fifth nucleotides, G41 and A42, in which G74 intercalates to form a continuous stacking interaction.

Other tertiary interactions important for stabilizing tRNA structure are also present in YrlA. For example, U32:A72 form a base pair equivalent to the Levitt base pair C48:G15 of tRNA^Phe^ (Figure 2C) (Levitt, 1969). This base pair is evolutionarily conserved in YrlA RNAs (Chen et al., 2014). In addition, in YrlA, U65-A71-A77, C70-G78-G30 and A66-U69-A79 form three base triplets that stack on one another. The equivalent base triplets are also found in tRNA^Phe^ (Figure 2D).

**Figure 2.**
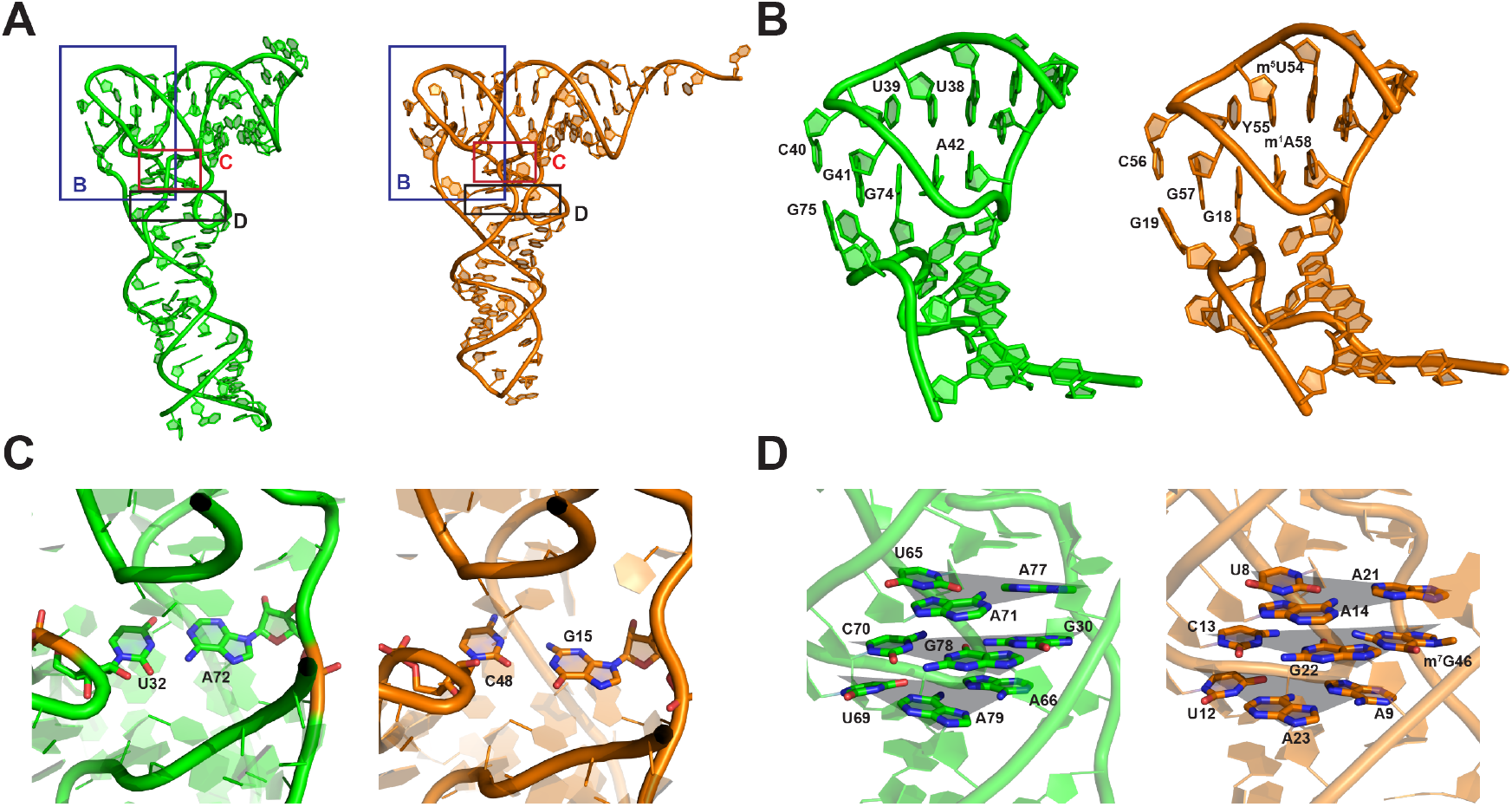
*S*. Typhimurium YrlA is stabilized by tRNA-like interactions. (A) Overall structures of *S*. Typhimurium YrlA (Green, PDB: 6cu1) and *S. cerevisiae* tRNA^Phe^ (Orange, PDB: 4tna). The areas shown in (B), (C) and (D) are boxed. (B) YrlA has DSL-TSL interactions that resemble that of tRNA^Phe^. (C) YrlA contains a Levitt base pair (A72-U32) similar to that of tRNA^Phe^ (G15-C48). (D) U65-A71-A77, C70-G78-G30 and A66-U69-A79 of YrlA closely resemble the base triples of tRNA^Phe^, U8-A14-A21, C13-G22-m^7^G46 and A9-U12-A23. The base triples are highlighted by shaded triangles (gray).

With the *S*. Typhimurium YrlA structure reported here, the crystal structures of three tRNA-like elements (TLE) have now been determined. The other two TLE structures are *Thermus thermophilus (T. thermophilus*) tmRNA and the TLS (tRNA-like structure) at the 3′-end of the turnip yellow mosaic virus (TYMV) genome (Bessho et al., 2007; Colussi et al., 2014; Gutmann et al., 2003). These structures share the L-shaped tRNA fold (Figure 3A) but not all tRNA features are present in all three TLEs. Notably, tmRNA and TYMV TLS contain free 3′-CCA ends and anticodon domain mimics (in the case of tmRNA, a portion of its SmpB protein partner substitutes for the anticodon stem-loop) (Bessho et al., 2007; Gutmann et al., 2003) (Figure 3A). The presence of these structural features is consistent with the ability of these two TLEs to be charged with amino acids and with their biological functions (Dreher, 2009; Keiler, 2015).

**Figure 3.**
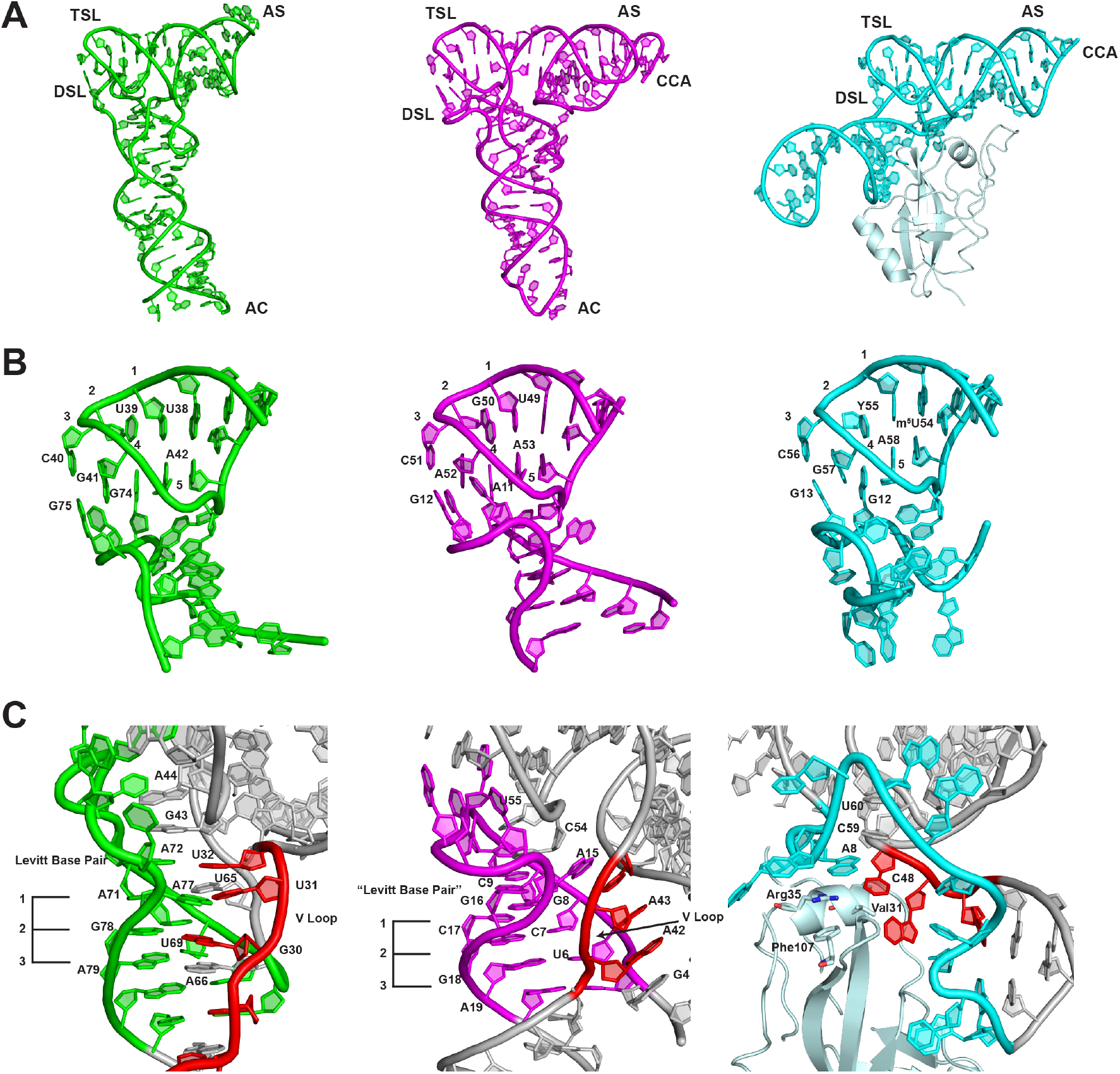
Comparison of the crystal structures of *S*. Typhimurium YrlA (Green, PDB: 6cu1), TYMV TLS (Magenta, PDB: 4p5j) and *T. thermophilus* tmRNA-SmpB complex (Cyan, PDB: 2czj). (A) The three TLEs share the same overall L-shaped fold. (B) The interactions between TSL and DSL are very similar between YrlA, TLS and tmRNA-SmpB. (C) The interactions between the VL and DSL show large differences but maintain the same architecture in the three TLEs. The VLs are colored in red. The three stacking layers of base triples or base pairs are labeled with numbers 1-3. [The numbering of TYMV TLS and *T. thermophilus* tmRNA-SmpB complex are based on previous publications (Bessho et al., 2007; Colussi et al., 2014)]

The DSL-TSL interaction is well conserved among the three TLEs (Figure 3B). This is perhaps not surprising as this is a major interaction defining the tRNA fold. In all cases, two guanine nucleotides in the DSL interact with a T loop through hydrogen bonding and base stacking interactions (Figure 3B). Since these three molecules have completely different evolutionary trajectories, the highly similar DSL-TSL interaction and hence the tRNA-like fold must be a key feature of their function, such that all three TLEs have either retained it (in the case of YrlA and tmRNA, which may have evolved from tRNA), or convergently evolved to acquire it (viral TLEs).

In contrast, the interactions between the DSL and the VL regions are quite different for the three TLEs. In YrlA, the nucleotides in the DSL region interact with the VL region and form three base-triplet layers, which are very similar to that of canonical tRNAs. These three base layers further stack with the Levitt base pair A72-U32 and with two nucleotides in the T loop, G43 and A44, stabilizing the L-shaped fold (Figure 3C). TYMV TLS uses a different strategy to stabilize the tRNA-like fold. Its VL does not interact with the DSL. Instead, all three residues in the VL (A42, A43 and U44) flip out to form a continuous stack with A3, G4 and A15. As a consequence, the nucleotides in the D stem form three base pairs instead of base triplets. Nonetheless, the DSL base pairs maintain a stacking interaction with two nucleotides in the T loop (C54 and U55) via a Levitt base pair equivalent (G16-C9) (Figure 3C). Strikingly, these interactions are also mimicked in the case of tmRNA but are conveyed through the SmpB protein, where Val 31, Arg35 and Phe107 bridge the base-pairing and stacking interactions. The completely different strategies used by the three TLEs to stabilize the VL-DSL connection highlight the importance of this region in maintaining the tRNA-like fold. These differences are also consistent with the fact that most tRNA-binding factors do not recognize this portion of tRNAs.

### The YrlA effector domain consists of a conserved tRNA core with variable stems and loops

A sequence alignment of YrlAs from various bacteria shows that the nucleotides involved in stabilizing the tertiary interactions are highly conserved (Figure 4A). These nucleotides include those in the D and T loops, the pyrimidine at the last position of the VL and the ^65^UA^66^ dinucleotide that connects the AS and the DSL. Most YrlA species also maintain the Levitt base pair (Levitt, 1969) that is important for stabilizing the L-shaped structure (^32^U and ^72^A in *S*. Typhimurium YrlA). Thus, we predict that YrlAs from other species will have the same overall tRNA-like fold as observed in the current structure.

**Figure 4.**
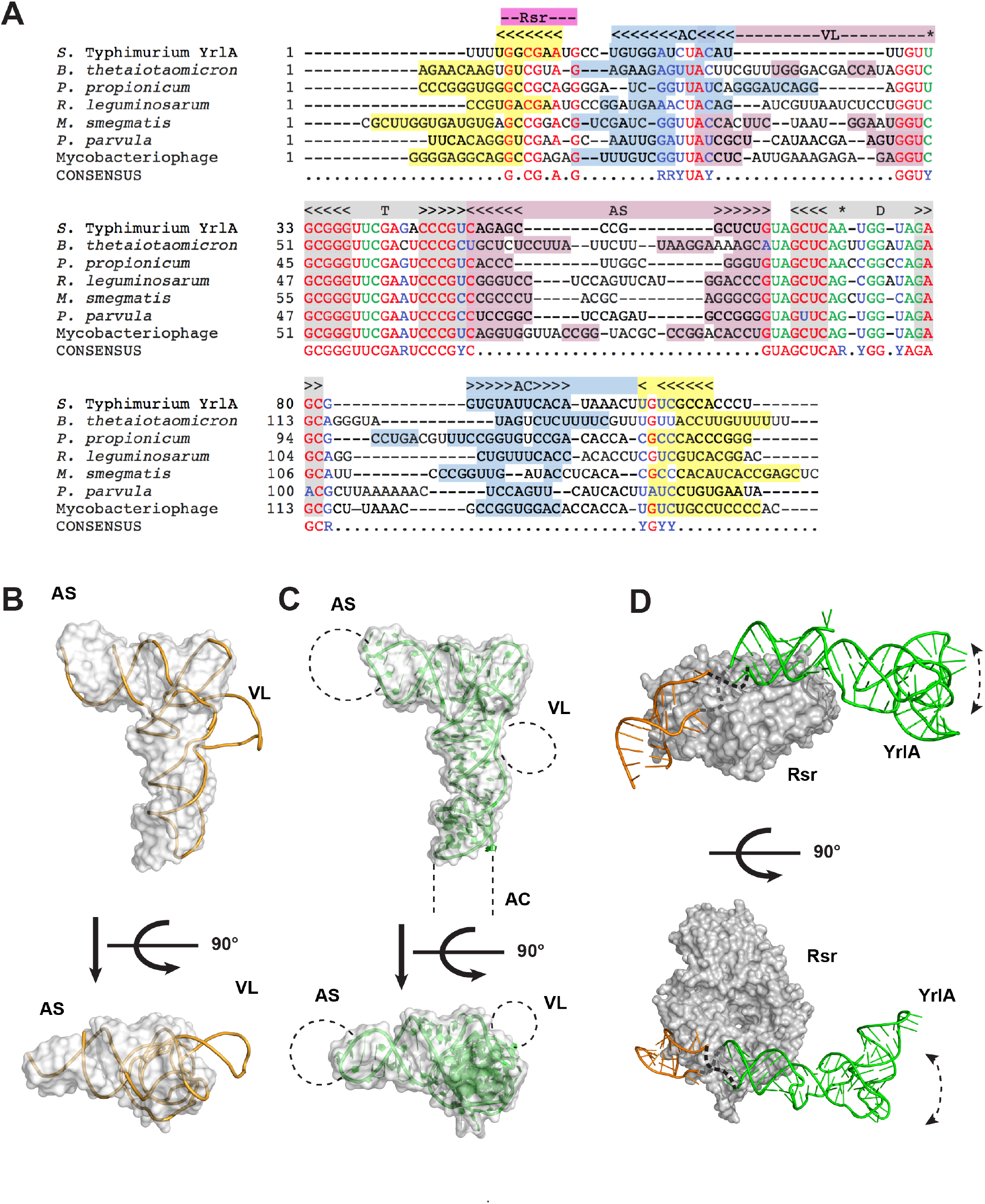
YrlA from various species likely share the same tRNA-like fold. (A) Sequences of representative YrlA RNAs were aligned using Clustal Omega (Goujon et al., 2010; Sievers et al., 2011) and further adjusted manually. The positions of the DSL and TSL are indicated in gray above the alignments and nucleotides that basepair to form the D- and T-stems are shaded gray. The positions of the AS and VL are indicated in pink, the AC is labeled light blue and the Rsr binding site is indicated with magenta. Nucleotides with the potential to form helices within the stem created by basepairing the 5′ and 3′ ends are highlighted light blue and yellow. Nucleotides conserved in YrlA RNAs are in red, with conserved nucleotides important for maintaining tertiary tRNA-like structure in green. Conserved nucleotides that are either purines or pyrimidines in YrlA RNAs are in blue. The shaded nucleotides in the VL region can potentially form duplexes. Nucleotides forming Levitt base pair are indicated with a star. (B) The structure of a representative class II tRNA, *T. thermophilus* tRNA^Tyr^ (1h3e, chain B, yellow) was superimposed onto the *S*. Typhimurium YrlA structure (gray surface). (C) Schematic model of a generalized YrlA RNA. The *S*. Typhimurium YrlA structure is shown as both cartoon (green) and gray surface. The different sizes of the AS, VL and AC regions are represented with dashed lines. (D) Model of the *S*. Typhimurium Rsr-YrlA complex. *X. laevis* Ro in complex with its Y RNA fragment (PDB: 1yvp) was used as a model for Rsr (gray surface) and the YrlA Rsr/Ro60 binding module (bright orange). The effector-binding module of YrlA is shown in green.

Although *S*. Typhimurium YrlA has a short VL, many YrlA species contain the long VLs (Figure 4A) that are characteristic of class II tRNAs. When a representative class II tRNA structure was superimposed on the structure of *S*. Typhimurium YrlA, only the VL protrudes from the L-shaped volume defined by this YrlA (Figure 4B). Class II tRNA VLs contain short duplexes, which could also exist in YrlA species with long VLs (Figure 4A). In addition, the sequence alignment indicates that the AS of YrlA RNAs vary in both sequence and length, as does the portion of the YrlA that corresponds to the AC. These comparisons support a model in which the YrlA core folds into a conserved tRNA-like structure, while the sizes of the AS, AC and VL regions vary (Figure 4C).

### Potential interactions of YrlA RNA with cellular factors

Interestingly, although sequences within the D and T stems are not required to maintain the tRNA L-shape, these nucleotides are conserved in YrlA RNAs (Figure 4A). This suggests that the YrlA D and T stems are under evolutionary selection pressure. Most aminoacyl-tRNA synthetases recognize the AC and the AS (Giegé and Eriani, 2014). In addition, the tRNA elbow, where the D and T loops interact, is recognized by the ribosome, ncRNAs and many tRNA-interacting proteins (Zhang and Ferré-D’Amaré, 2016). To our knowledge, no cellular factors have been shown to function by recognizing specific sequences within the T and D stems of tRNAs or TLEs. The sequence conservation in this region suggests that YrlA RNAs may interact with a novel factor that recognizes specific sequences in these stems.

We compared available structural information to gain insight into the possible architecture of the Rsr-YrlA. YrlA RNAs contain a conserved Rsr/Ro60-binding module (Figure 1A); hence the structure of the Rsr-YrlA module is believed to resemble that of the *X. laevis* Ro60-Y RNA complex (PDB: 1yvp) (Stein et al., 2005). For *S*. Typhimurium YrlA, the tRNA-like effector-binding module, with structure reported herein, is connected to the Rsr/Ro60-binding module by unpaired short loops of 5 nucleotides. This allows us to model the Rsr-YrlA complex by positioning our structure close to the upper surface of the *X. laevis* Ro60-Y RNA structure (Figure 4D, upper panel). This surface of Ro60 presents positive charges that were shown to be important for both Y RNA and misfolded 5S rRNA binding (Fuchs et al., 2006; Stein et al., 2005). Thus, as predicted for vertebrate Y RNAs and Ro60, YrlA could serve as a gatekeeper to regulate access of other RNAs to Rsr.

The unknown effector factor that binds YrlA is presumably bound to the elbow region of the YrlA tRNA-like domain, more specifically, the T stem and the D stem. In the presence of misfolded RNA and/or the effector protein, the tRNA-like domain of YrlA may reposition. For instance, the architecture of the *D. radiodurans* RYPER has been determined by single particle electron microscopy (EM) and the Y RNA is predicted to bend downward from the Rsr surface, allowing the stemloop-containing module to contact one or more S1/KH domains of PNPase (Chen et al., 2013). The architecture of the *S*. Typhimurium Rsr-YrlA-effector complex remains to be determined.

## DISCUSSION

Although the structure of the metazoan Y RNA module that binds the Ro60 autoantigen was elucidated using X-ray crystallography (Stein et al., 2005), high resolution structures of the effector-binding domains of these RNAs have been lacking. Our crystal structure of the tRNA-like module of *S*. Typhimurium YrlA RNA reveals that this module not only adopts an overall L-shaped structure similar to tRNA, but is also stabilized by the same tertiary interactions. Since all sequences involved in critical tertiary interactions are strongly conserved in YrlA RNAs, we predict that the ability to fold into the canonical tRNA L-shape is a general feature of this Y RNA family. Moreover, the high degree of sequence conservation at the YrlA region corresponding to the tRNA elbow, particularly within the T and D stems, contrasts with the variable sizes of the AS, AC and VL regions. Based on the extreme conservation of these T and D stem sequences, we expect that the effector(s) that bind YrlA RNAs will be one or more tRNA-binding protein(s) that recognize the elbow region with some sequence specificity.

The strong resemblance of YrlA RNAs to tRNAs lends support to the proposal that YrlA and other Y RNAs evolved from tRNA (Chen et al., 2014). Consistent with this hypothesis, Y RNAs are encoded adjacent to one or more tRNAs in some bacteria (Chen et al., 2014). Additionally, the finding that YrlA RNAs differ from *bona fide* tRNAs in that the TSL occurs 5′ to the DSL supports a recent model in which these RNAs originated from dimeric tRNA transcripts (Sim and Wolin, 2018). In this model, the YrlA TSL derived from the first tRNA, while the DSL derived from the second tRNA. Since the YrlA AS would originate from the spacer between the two tRNAs, a model in which YrlA evolved multiple times in distinct bacteria would provide an explanation for the variable length of this stem. Alternatively, if YrlA evolved from a single primordial dimeric tRNA, there may have been less pressure to maintain the length of the AS. In either case, the additional sequences in a dimeric pre-tRNA, such as the DSL and AC of the first tRNA and the AC and TSL of the second tRNA, could potentially basepair to form a stem containing a sequence recognized by Ro60.

In certain algae and at least one archaeal species, some tRNAs are transcribed as circularly permuted variants that are processed to mature tRNAs (Chan et al., 2011; Maruyama et al., 2010; Soma et al., 2007). Some of these pre-tRNAs resemble YrlA in that the AS is initially a closed loop and the AC stem is initially formed by base pairing the 5′ and 3′ ends of the newly made RNA. These unusual pre-tRNAs are processed to canonical tRNAs by excising and ligating the extended AC stem to form a circular intermediate, followed by opening of the AS by endonucleases such as RNase P and/or RNase Z (Soma et al., 2007). Although enzymes equivalent to the eukaryotic and archaeal splicing endonucleases have not been reported in bacteria, the resemblance of YrlA RNAs to circularly permuted tRNAs raises the possibility that some YrlA RNAs could undergo processing to more closely resemble canonical tRNAs.

The tRNA resemblance is less evident for Yrn1 RNAs: however, these ncRNAs also contain some tRNA-like features. Yrn1 can be folded to contain a TSL that conserves the T stem sequences of YrlA RNAs (Chen et al., 2014). This TSL likely forms in vivo, as it contains pseudouridine at the position corresponding to the pseudouridine in TSLs of all canonical tRNAs (Chen et al., 2014). Since structures have not been reported for the effector-binding domain of Yrn1 or any metazoan Y RNAs, it remains possible that the three stem loops in the Yrn1 effector-binding domain fold in three-dimensions to mimic tRNA.

Although the exact role of YrlA RNA is unknown, it is likely that it functions in RNA degradation and/or repair. Consistent with a role in RNA degradation, some YrlA and Rsr co-purify with PNPase in S. Typhimurium (Chen et al., 2013). If, as described for Yrn1 (Chen et al., 2013), the YrlA effector-binding domain interacts with the S1 and KH domains of PNPase, the highly folded tRNA domain could serve to protect the RNA from endonucleolytic nicks that would render it a substrate for PNPase or other exoribonucleases.

Rsr, YrlB and YrlA have also been proposed to function in RNA repair, since they are encoded adjacent to the RtcB RNA ligase in many bacteria (Burroughs and Aravind, 2016; Chen et al., 2013; Das and Shuman, 2013). In some bacteria, including *S*. Typhimurium, this operon (*rsr-yrlB -rtcBA*) encodes both RtcB, the ligase that joins pre-tRNA halves following intron excision in Archaea and metazoans (Englert et al., 2011; Popow et al., 2011; Tanaka et al., 2011; Tanaka and Shuman, 2011), and RtcA, an RNA terminal phosphate cyclase (Das and Shuman, 2013; Filipowicz et al., 1985). Although the substrates of RtcB in bacteria are largely unknown, *E. coli* RtcB repairs 16S rRNA following cleavage by the MazF toxin (Temmel et al., 2017). Because Rsr and Y RNAs are encoded adjacent to RtcB in bacteria from multiple phyla, it was proposed that Rsr and one or more Y RNAs function as cofactors to enhance RtcB activity (Burroughs and Aravind, 2016). Consistent with a more general role in RNA ligation, Rsr and YrlA are occasionally encoded adjacent to members of other RNA ligase families.

Interestingly, in certain other bacteria, RtcB is encoded adjacent to a protein containing a Band-7 domain and a predicted ncRNA, called band 7-associated tRNA (b7a-tRNA), that strongly resembles an authentic tRNA (Burroughs and Aravind, 2016). Consistent with functional redundancy between b7a-tRNA and YrlA, these bioinformatics searches predict that occasionally b7a-tRNA is encoded adjacent to Rsr and YrlA is adjacent to the Band-7 domain protein. Although the existence of the putative b7a-tRNA has not been validated experimentally, these predictions, together with our finding that the YrlA effector-binding domain folds similarly to tRNA, support the hypothesis that tRNA-like molecule(s) contribute, directly or indirectly, to RNA ligation.

## MATERIALS AND METHODS

### Plasmid construction and RNA purification

The tRNA-like domain of *S*. Typhimurium YrlA was cloned into the EcoRI and NheI sites of plasmid pHDV4 (Walker et al., 2003), such that the YrlA coding sequence was followed by the HDV ribozyme. *S*. Typhimurium YrlA was transcribed from HindIII-linearized plasmid using T7 RNA polymerase (Milligan et al., 1987). The transcription reaction was mixed with an equal volume of ribozyme denaturing buffer (8M Urea and 0.5M MgCl_2_) and incubated at 37 °C for 2h to increase HDV ribozyme cleavage efficiency (Rosenstein and Been, 1990). Following T7 transcription and HDV cleavage, the sequence of the resulting RNA is: <u>GG</u>GUGG<u>C</u>UCUACAUUUGUUGCGGGUUCGAGACCCGUCAGAGCCCGGCUCUGUAGCUCAAUGGUAGAGCGGU<u>G</u>UAGUCAC<u>C</u>. The underlined nucleotides indicate modifications to the original YrlA sequence in order to stabilize the AC stem. The YrlA product was precipitated with ethanol and purified by polyacrylamide-urea gel electrophoresis. YrlA variants were made using QuikChange Site-Directed Mutagenesis (Agilent).

### Crystallization and data collection

The purified YrlA RNAs were folded by heating to 95 °C for 2 min, transferring to 60 °C and incubating for 2 min. MgCl_2_ was added at 60°C to a final concentration of 10 mM and the reaction was quenched on ice for 30 min before use. The folded RNA was then buffer exchanged to RNA crystallization buffer (50mM sodium cacodylate, 50 mM KCl, 1 mM MgCl2 and 0.1 mM EDTA) using Amicon Ultra Centrifugal Unit (Merck Millipore, Billerica, MA). The RNA was concentrated to a final concentration of 1.2-1.5 mg/ml and screened for crystals using the microbatch under oil method using the Nucleix Suite (Qiagen, Germantown, MD). Crystals were readily formed under numerous conditions overnight.

Three YrlA variants were crystallized, namely YrlA WT, YrlA Tetraloop and YrlA Tetraloop 3C, in which an additional cytosine was added to the 3′-end of YrlA Tetraloop. The best crystallization conditions for each YrlA constructs were summarized in Table S1. Crystals were cryoprotected by Paratone oil (Hampton Research, Aliso Viejo, CA) or 30% glycerol. Diffraction data was collected at the Advanced Photon Source beamlines 24ID-C and 24ID-E. The data statistics are summarized in Table 1.

### Structure determination and refinement

The structure of YrlA RNA was determined by a combination of single wavelength anomalous dispersion (SAD) and molecular replacement. The initial phase information was obtained by SAD phasing using SHELX C/D/E (Sheldrick, 2008) from a data set collected on an Iridium hexamine derivative of the YrlA-Tetraloop-3C crystal. A preliminary model was built and used as the search model for the YrlA-Tetraloop data set using Phaser (McCoy et al., 2007). There is one RNA molecule in the asymmetric unit of the crystal. The initial model was refined by iterative rounds of restrained refinement using refmac5 (Vagin et al., 2004) followed by manual rebuilding with Coot (Emsley and Cowtan, 2004). B-factor sharpening was performed to facilitate model building (Liu and Xiong, 2014). The structure was refined to final R/R-free of 21.3%/23.4% with excellent electron density (Figure S1). Refinement statistics are summarized in Table 1.

## ACKNOWLEDGEMENTS

The authors would like to thank the staff at the Advanced Photon Source beamline 24-ID for assistance in data collection. This work was supported by National Institutes of Health Grants R01AI116313 (Y.X.) and R01GM073863 (to S.L.W.) and by the Intramural Research Program, Center for Cancer Research, National Cancer Institute, National Institutes of Health (X.C. and S.L.W.).

The authors declare no conflicts of interest.

## AUTHOR CONTRIBUTIONS

X.C., S.W., and Y.X. conceived the project. W.W., X.C., S.W. and Y.X. designed the experiments. W.W. and X.C. purified the YrlA RNAs and W.W. crystalized the RNA and collected the X-ray diffraction data. W.W. and Y.X. determined the structure. W.W., X.C., S.W. and Y.X. analyzed the data and prepared the manuscript.

